# Different phylogenetic characteristics of Icterids: would they be evolutionary strategies to avoid parasitism by Molothrus spp.?

**DOI:** 10.1101/2024.06.30.601453

**Authors:** Vinícius Munhoz Barbosa, Bianca Dinis da Silva, Vinícius Xavier da Silva, Érica Hasui

## Abstract

Some birds exhibit the behavior of nest parasitism, which involves laying their eggs in the nests of other species to be incubated and cared for by the adoptive parents. Among all the studies conducted on this subject, there is a gap regarding the nest type of nest parasites and their hosts. Therefore, using species from the Icteridae family, this study aimed to identify if there is a tendency for more closed nests to be less parasitized than open nests and if there is a phylogenetic relationship between them. In this context, we expected open nests to be an ancestral condition to closed nests, serving as an evolutionary feature to avoid nest parasitism. We also analyzed other characteristics such as the number of eggs, nest type and parental care. As a result, we observed that open nests were more common, while closed nests were predominant in a specific clade and some isolated species. The analyses indicated a phylogenetic signal clustered within the Icteridae family concerning nest types, which may imply a selective pressure. However, we cannot assert that it is a direct response to nest parasitism, as closed nests are also parasitized, specifically by *M. oryzivorus*. Parental care and diet type also showed phylogenetic signal, indicating that these changes were not random. However, we did not observe associations in host selection by the parasites based on these characteristics. Furthermore, we found a progression in the number of species parasitized by *Molothrus* spp. along the phylogenetic lineage. We also observed a similarity in host choice between *M. ater* and *M. aeneus*, indicating evolutionary convergence, as they are not sister groups.

## INTRODUCTION

The brood parasitism is the behavior observed in some birds that don’t build their own nests and lay their eggs in the nests of other species to be incubated and raised [1]. For the host, taking care of a parasitic chick demands higher energy costs and may also lead to lower survival rates for its own chicks [2]. This activity becomes even more challenging because the parasitic birds can drill into the eggs or kick out the host’s chicks from the nests to reduce competition for food [3, 4].

Due to these energy and reproductive costs, evolutionary strategic responses are expected in hosts to avoid nest parasitism. Likewise, responses from parasites are anticipated in order to overcome the strategies imposed by the hosts and successfully engage in nest parasitism. This coevolutionary mechanism between parasites and hosts is known as an arms race [5].

In a scenario that considers only the parasite-host interaction without the occurrence of an arms race, species with phenotypic disadvantages would experience a decrease in individuals. In the context where the arms race naturally takes place, less common phenotypes may end up prevailing over more common ones, thereby altering the current outcomes of the “war” [6]. Thus, both parties undergo this process through evolutionary feedback to overcome the phenotypes developed by their opponents, generating a cycle of evolutionary changes in the parasite-host interaction to gain advantages against new evolutionary strategies.

One of the strategies developed by hosts is the recognition of their own eggs [7] and consequently, the rejection of parasitic eggs [8]. Within this context, a population of a host species may exhibit eggs with different morphological attributes compared to a population that is not affected by parasitism [9]. Moreover, among studies that analyze evolutionary mechanisms to avoid parasitism, LÓPEZ et al. (2021) found an increase in shell resistance in the eggs of host birds, making it difficult for parasites such as *M. rufoaxillaris* and *M. bonariensis* to penetrate them. In host species of *M. ater*, the same study found larger and more conical eggs, which hinder their removal by the parasite.

On the other hand, the selection process in parasites is focused on increasing the similarities of the eggs to those of the hosts [11]. In addition to egg characteristics, several strategies can maximize the survival of parasite offspring. For example, MORELLI, BENEDETTI, and PAPE MØLLER (2020) found that parasitic cuckoo species have a more specialized diet compared to non-parasitic species. Cuckoos that develop in nests of seed-feeding species, for instance, are likely to perish [13] since most cuckoos have an insect and invertebrate-based diet [13]. These differences can hinder the survival of parasite offspring as the hosts may not provide a suitable diet for them. This raises the question of whether the host’s diet is used as a selection criterion by the hosts.

We further assume that some factors not yet mentioned can influence the success of parasitism, such as parental care and the number of eggs laid. In this regard, parental care is essential for the development and protection of the host’s own offspring. When both parents provide care, it can be considered more advantageous for the host than when care is provided by only one parent, as they share tasks and invest time and energy in caring for an extra chick [2], which also increases the chances of survival for the host’s offspring. Lastly, the number of eggs also has an influence, as a large quantity of eggs would lead to increased competition for food within the nest and require the parents to spend more time outside the nest in search of food.

Based on the studies reviewed by MEDINA et al. (2020), we obtained a general overview of what has been discussed. However, according to the study, little consideration has been given to the possible interaction between the evolution of host nest structures and the presence or absence of parasitism in species. We have found only studies that focus on one aspect or another of what we want to investigate. For example, the study by ANTONSON et al. (2020) discusses the protection provided by closed nests, which can offer thermoregulatory benefits, but it does not specifically address the evolutionary aspect of nests. There are also studies on nest structure as an evolutionary form of defensive effectiveness [2, 17, 18], but they also do not explore the evolutionary history of nest formats throughout the phylogeny.

The study by RUTILA et al. (2002) demonstrates that cavity nests reduce the success of cuckoo egg laying due to the large size of the parasitic species. In addition, weaverbirds construct nests with long entrance tubes that act as protection against nest parasitism and predation [2, 17], with records of parasitic cuckoos getting trapped when attempting to access the nest [2]. Therefore, we aim to connect the information in order to elucidate the evolutionary history along with the parasite-host interaction.

With this in mind, our objectives were: (I) To investigate the possible correlation between different host nest formats, diet, parental care, and number of eggs in the choice made by parasites. (II) To evaluate the phylogenetic relationship between nest formats, examining whether there is an evolutionary trend towards increasingly closed nest conditions as an anti-parasitic adaptation. (III) To analyze whether there is a progressive increase in the number of parasitized species throughout the evolutionary process of Molothrus. (IV) To investigate if there is similarity in host choice by parasites and whether this similarity is related to the phylogenetic proximity between the parasites.

We expect to find an evolutionary trend where open nests are more basal in the phylogeny, and closed nests with corridors are more recent, representing an adaptive response of hosts to the pressure from parasites (FIG 1). Regarding diet, our expectation is that parasites select hosts with a similar diet to their own. With the exception of *M. aeneus*, all other species have hosts that feed on vertebrates, while *M. aeneus* hosts feed on seeds. This factor may be relevant for feeding success, as in the case of cuckoos, the nest parasites have a different diet compared to non-parasitic species [12].

**Fig 1.**
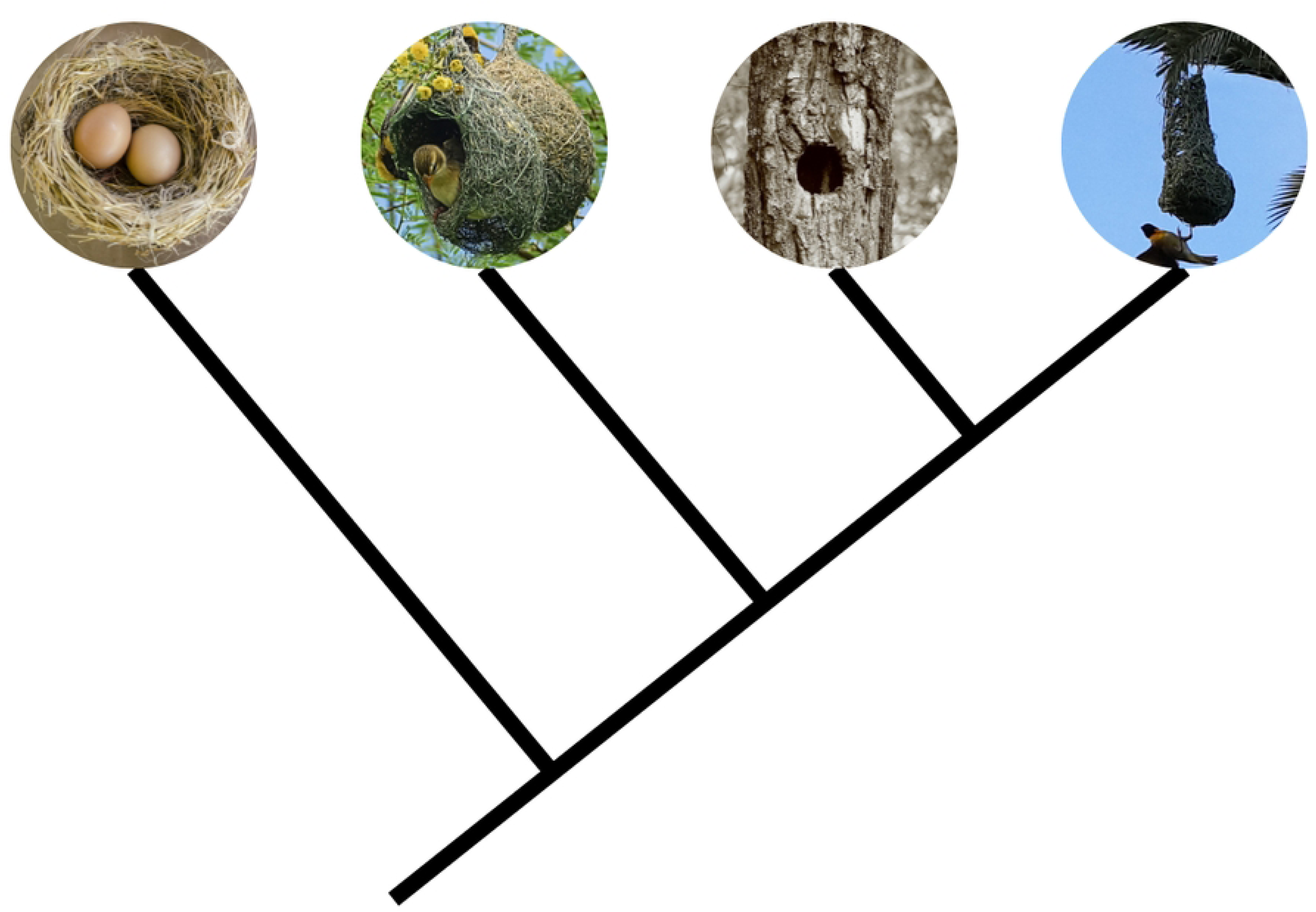
Cladogram depicting the evolutionary hypothesis of nest types. Phylogenetic hypothesis of nest type evolution, where the most basal type is open nest, leading to the most recent evolutionary form of a closed nest with a corridor. Reproduced by the authors based on Crozariol, M. A, 2016.

Regarding the number of eggs, our expectation is to find a smaller clutch size due to the effort of host parents in providing food. More offspring would result in reduced parental investment per individual due to increased costs, leading to a lower feeding rate and/or competition among the chicks [20], although other factors such as resource abundance may also be associated In terms of parental care, we expect a preference for hosts where only the female provides care because although two parents may be able to feed the parasites and/or protect the nest better, the vigilance against invaders is also important. For example, there is an evolutionary strategy where birds form reproductive coalitions, with multiple birds of the same species nesting in close proximity to each other as a response to the pressure from nest parasites, aiming to hinder the success of parasitic egg laying [21], even though successful invaders may encounter increased protection.

Finally, we may also find a growing trend of species parasitized by the genus *Molothrus*, as a result of the evolutionary success of this strategy, similar to the phylogeny proposed by LANYON (1992) that includes all parasitized species across different families.

## Materials and methods

### Target species of study

We selected the species of Icteridae that have at least one record of brood parasitism by one or more species of the genus *Molothrus*, namely *M. bonariensis*, *M. aeneus*, *M. ater*, *M. rufoaxillaris*, and *M. oryzivorus*, based on the classification by LOWTHER (2018). *M. rufoaxillaris* and *M. oryzivorus* are brood parasites of a smaller number of host species compared to the others. The former parasitizes 5 host species of icterids, while the latter parasitizes 9 icterids and 2 corvids, defining them as specialists in comparison to *M. aeneus*, which parasitizes 16 families, *M. bonariensis*, which parasitizes 27 families, and *M. ater*, which parasitizes 40 families, thus considered generalists [23].

Among all 11 families, the Icteridae family has the highest species richness parasitized by *Molothrus* spp. in the Americas (103 species), and it is also the only family that is parasitized by all five brood parasite species [23], making it the chosen family for our study.

### Level of brood parasitism

Based on the classification by LOWTHER (2018), we generalized the categories “host” for cases of successful brood parasitism and “victim” for cases of unsuccessful parasitism. “Victim” represents species that had parasitic eggs placed in their nests, but with empirical evidence of failure in the development of the parasitic offspring. On the other hand, the term “host” represents species that had parasitic eggs placed in their nests, with successful parasitism taking place [23]. The remaining species are either not parasitized by the genus *Molothrus* or simply do not co-occur with its species, which, for practical purposes of the analysis, has the same meaning.

### Nest type

The categorization of nest types was based on the standardization of SIMON and PACHECO (2005). The external nest shape was disregarded unless it interfered with the internal structure. Therefore, nests that, for example, are open and have a thicker or thinner wall do not differ in terms of the parasite’s access to the nest.

Thus, the nests were categorized into four categories: (1) open (cup-shaped); (2) semi-closed; (3) closed (such as the nest of the *Rufous hornero* or a tree cavity); and (4) closed with access corridor (pouch-shaped) (Fig 1).

### Collected data

In addition to the list of species parasitized by each *Molothrus* species, we collected data on five other characteristics of these icterids’ biology that could be related to more intense or less intense nest parasitism: nest type, diet, parental care, number of eggs (Supporting information: Table 1). To gather the data, we compiled information from various bibliographic sources available on digital platforms and citizen science platforms such as eBird (http://www.ebird.org), WikiAves (www.wikiaves.com.br), Animal Diversity Web (ADW) (https://animaldiversity.org), and Birds of World (https://birdsoftheworld.org/). Nest types were classified based on images or descriptions. In cases where the websites provided different quantities for the number of eggs of each species, we calculated the average of the values presented. We also obtained information on hatchling mass, adult mass, and diet from SHEARD et al. (2019).

### Co-occurrence analysis

From the parasitic species, we analyzed their co-occurrence with each icterid species using the geographic distribution maps of each species [26]. This analysis was necessary because co-occurrence associated with non-parasitism could contain relevant evolutionary or adaptive information, thus testing hypothesis II.

For this analysis, we utilized ArcMap software v. 10.8 [27] to read the data. We strictly considered as co-occurrence only the species that had some degree of overlapping distribution with the *Molothrus* spp. In cases where species were not present in the RIDGELY et al. (2007) files, we used data from the IUCN Red List Threatened Species (Supporting information: Table 2).

### Definition of bioregions

The division into bioregions was necessary to assist in the understanding of the parasite-host interaction. Therefore, we developed the bioregions with the aim of achieving a balanced distribution of samples and ensuring sufficient and well-distributed sampling capable of representing the co-occurrences of parasites and hosts, as these environments are characterized by unique sets of temperature variation, topography, among others [28].

The bioregions were defined for statistical analysis using the cluster technique, where we selected samples of icterid assemblages based on similarities in species composition. For this purpose, we used the IUCN website to search for all icterid species in the Americas and downloaded their distribution data in the form of “data” files (Supporting information: Data_0). With this data, we created the bioregions using Infomap Bioregions, a web application network for identifying biogeographic regions [29].

Infomap Bioregions is an integrated data network capable of generating bioregions based on polygonal and/or point data of species distributions [29]. Depending on the amount of expressed data, the distribution grids can be larger when there are few data points or smaller when there are many. In this way, it is possible to generate bioregions that are comparable to traditional standardized classifications [30, 31], such as continental barriers that prevent species migration between continents. With the help of the application, we obtained 22 bioregions (Supporting information: Bioregion and Table 2).

### Sample point selection

We selected 8 out of the 22 bioregions, as they had the most complete data for obtaining the sample points, from which the species present were then organized into a new spreadsheet (Supporting information: Table 3). This selection of sample points was necessary for conducting statistical calculations based on cooccurrence relationships.

From this, we created six random sample points in each bioregion with a minimum distance of 100 km between them to avoid overlapping community occurrences, using QGis software v. 3.26.2 [32]. In bioregions that were too small to include six points, we opted to use all available points. Finally, we created a list with all the species present (Supporting information: Table 3).

### Phylogenetic signal and ancestral state reconstruction

We analyzed the phylogenetic relationships to understand how the studied traits behave throughout the evolution of Icteridae species. We obtained 1000 phylogenies for the regional species pool extracted from JETZ et al. (2012), which were obtained from the birdtree website (http://birdtree.org/), and constructed a majority-rule consensus cladogram using the software Mesquite v. 3.70 [34] (Supporting information: Consensus cladogram). This consensus cladogram served as the basis for subsequent analyses.

We analyzed the phylogenetic signal of the studied traits (Section 2.4), which indicates whether certain states of a trait are conserved throughout a phylogeny, meaning that closely related species exhibit similar states of variation for a particular trait. For example, the closed nest with corridor state of the nest type trait is present in two sister genera of Icteridae (*Psarocolius* spp. + *Cacicus* spp.). In contrast, we have what is known as convergent trait, where the same state of variation for a particular trait appears independently in distantly related species in the phylogeny (trait or state without phylogenetic signal) [35].

To assess the phylogenetic signal of the examined traits, we used the Picante Package in R platform v. 3.5.3 [36] with the “multiPhylosignal” command, based on the K statistic [37, 38]. The Blomberg’s K statistic is guided by the Brownian motion evolutionary model and can assume values between zero and infinity. When it does not differ statistically from zero, it indicates that species are randomly similar in a given trait (or that the trait has no phylogenetic signal or is convergent). When K differs from zero (p<0.05), it means that closely related species are, on average, more similar to each other in that trait (or that the trait has a phylogenetic signal or is conserved). When K approaches 1, there is a strong phylogenetic signal according to the Brownian motion model, and when much higher than 1, K indicates that the Brownian motion is not the best evolutionary model to interpret the evolution of that trait [39].

We reconstructed the ancestral state of each analyzed trait using the parsimony method because the maximum likelihood method can yield inconsistent results due to the influence of branch lengths in the phylogeny [40–42]. Characterizing the ancestral state of each trait allows us to understand which states of the traits were more conserved over time and how they have changed, thus identifying which evolutionary aspects have been retained and where new aspects have emerged. For this purpose, we used the Mesquite software v. 3.70 [34].

### Phylogenetic correlation

The Mesquite software allowed us to mirror (Mirror command) the phylogeny of the regional species pool, comparing the different states of the studied characteristics to test the phylogenetic correlation between them using the Pairwise Comparison command. This test is not a traditional statistical correlation precisely because it considers the phylogenetic relationships, as the characters of each species are not independent due to their varying degrees of common ancestry [34]. This correlation tests the association between all possible pairs of closely related species that differ in the states of two characteristics (let’s say characteristic 1, with states A and a, and characteristic 2, with states B and b). The test assesses the probability of the state A (or a) of one species in the pair being associated with the state B (or b) of the other species. It does not analyze closely related species with the same state for a particular characteristic, precisely because there is a high chance that this situation is dependent on common ancestry. However, although it reduces the chance of phylogenetic dependence, this algorithm does not guarantee 100% phylogenetic independence [43].

We also tested the correlation between all possible pairs of the characteristics investigated, particularly those that showed phylogenetic signal, and we also tested potential associations involving characters without signal. We analyzed these characteristics without apparent signal because even if they are not supported by a single lineage set in the phylogeny of the regional species pool, it does not mean that they cannot occur in other evolutionary scenarios or in analyses at a more restricted geographic scale where not all species from the regional pool are present [44]. We also conducted correlation analyses within the selected bioregions to consider real co-occurrence conditions. Additionally, even if a characteristic is shown to be convergent (emerging in independent lineages), it can still be an adaptive response to brood parasitism; it just may not be supported in phylogenetic terms [44].

## Results

### Phylogenetic signal

#### Nest types

From the consensus phylogeny, we found that the nest type exhibits significant phylogenetic signal in icterids (K = 0.1410, p = 0.001, Fig 2). This indicates that evolutionary changes are not random. The most basal condition observed was the open nest. Semi-open and closed nests have evolved multiple times independently throughout the family’s evolution, suggesting convergent evolution. Nest type is not correlated with parasitism, as we found hosts with both open and closed nests.

**Fig 2.**
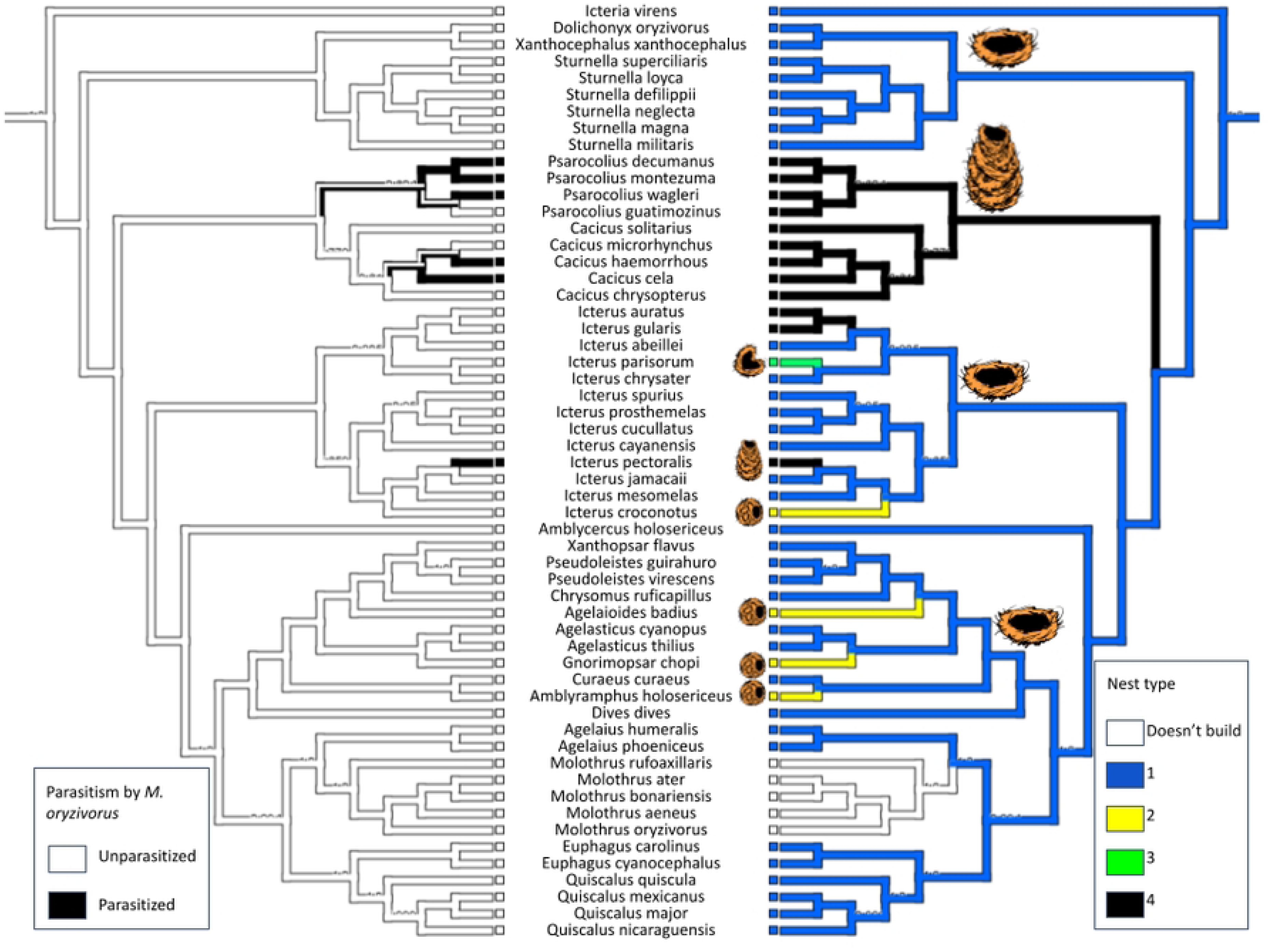
Parasitism by *M. oryzivorus* and nest types made by Icteridae species. The left side shows the species parasitized by *M. oryzivorus*. In black, we have the parasitized species, while in white, non-parasitized species. The right side displays the nest types of Icteridae species. Value 0, in white, represents *Molothrus* species that do not build their nests. Value 1, in blue, represents Icteridae species with open nests. Value 2, in green, represents semi-open nests. Value 3, in yellow, represents closed nests. Value 4, in black, represents closed nests with a corridor. Authors’ production.

*M. oryzivorus* showed a clear preference for closed nests of the genera *Psarocolius* and *Cacicus* (Fig 2). However, the correlation was not significant, as several species within these genera were not parasitized (p > 0.5). This indicates that nest condition alone is not a determining factor for parasitism in this species.

#### Diet types

Our statistical calculations demonstrated that there is a phylogenetic signal for diet type (k = 0.080, p = 0.036), once again indicating a non-random evolution. However, the states of the characteristics are widely spread among carnivores, omnivores, and invertebrate feeders, with the latter being the most basal (Fig 3).

**Fig 3.**
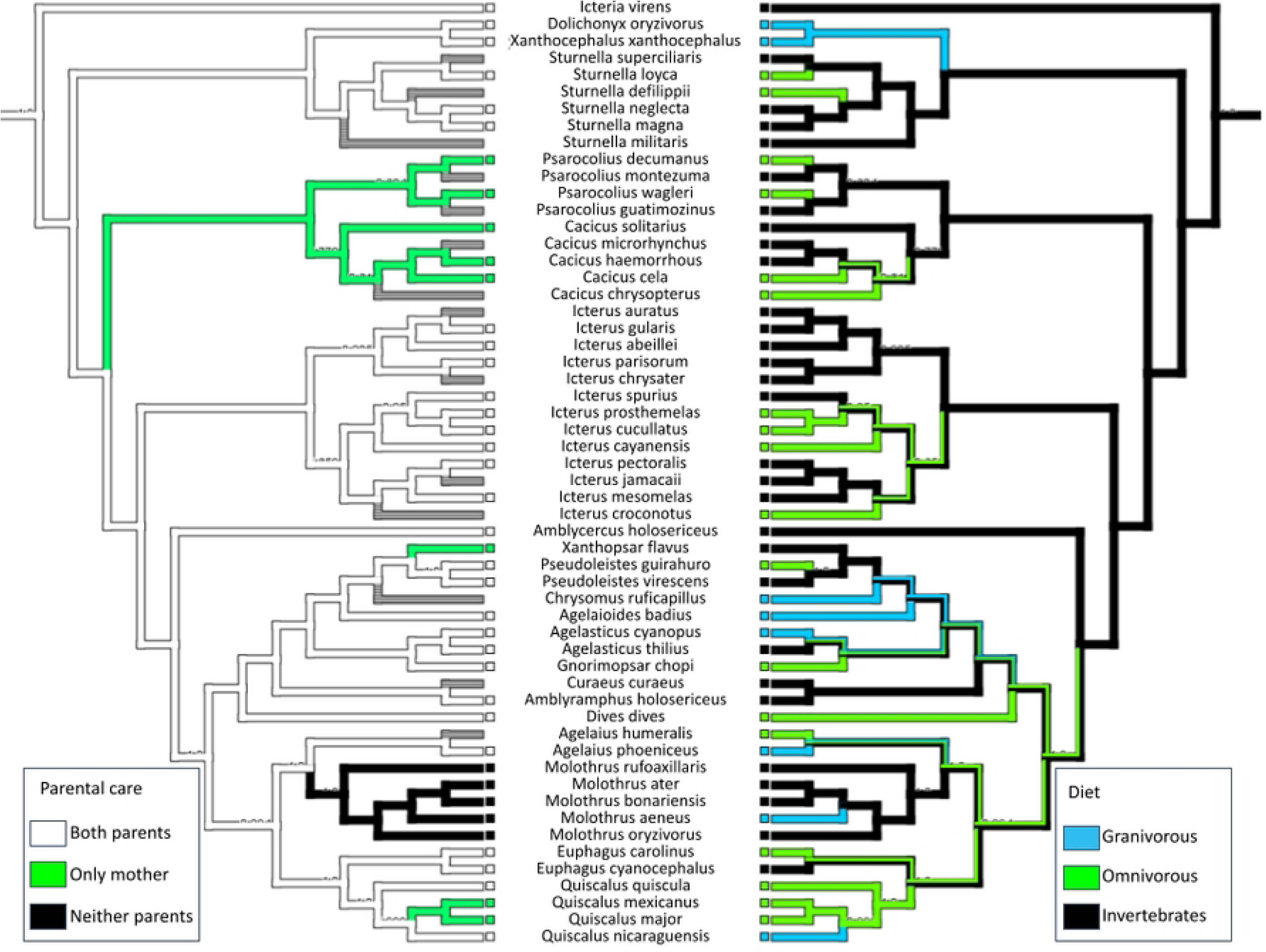
Correlation between Parental Care and Diet in Icterids. In parental care (on the left), the white color indicates that both parents provide care, the green color indicates that only the mother provides care, and the black color indicates that neither parent provides care (brood parasites). In terms of diet (on the right), the blue color represents seed-based diets, the green color represents omnivorous diets, and the black color represents diets based on invertebrates. Authors’ production.

#### Parental Care Types

Parental care also showed a phylogenetic signal (K = 0.3885, p = 0.001). The most basal state is that both parents care for their offspring, and throughout the clade, there are branches where only mothers provide care, as well as the branch of the brood parasites themselves, which do not engage in parental care (Fig 3).

### Phylogenetic correlation tests

#### Phylogenetic test for regional pool

Among the analyses conducted with the regional pool, the correlation between *M. ater* and the number of eggs was marginally significant (p = 0.0625), indicating its preference for nests with more than 4 eggs (Fig 4). The marginal result was obtained because *M. ater* also included some host species with 3 and 4 eggs in its parasitism. Regarding the similarity in host selection by the parasites, we observed a significant correlation between *Molothrus aeneus* and *Molothrus ater* (p=0.015625), indicating that the two species parasitize many species in common (Fig 5).

**Fig 4.**
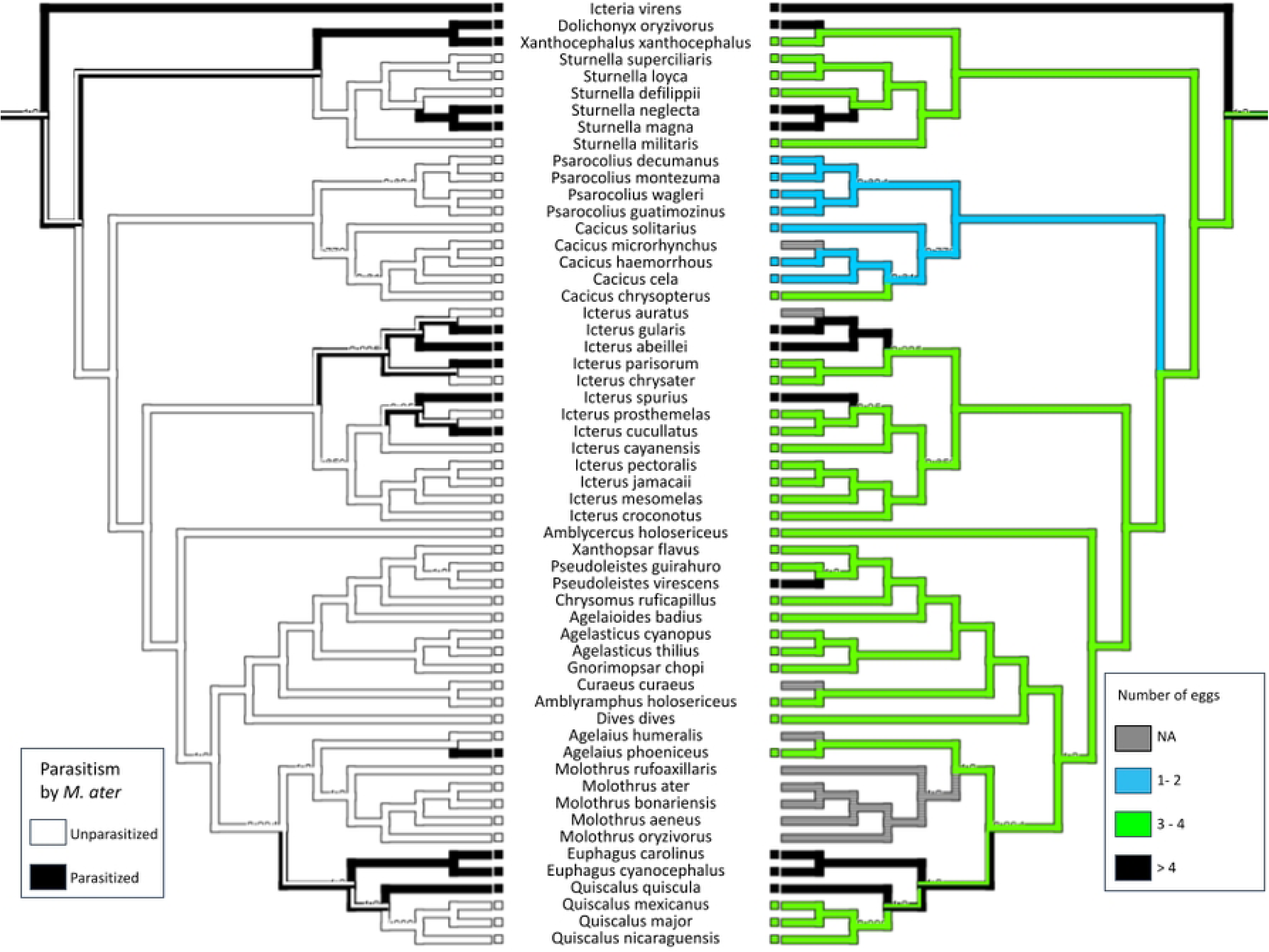
Correlation between the parasitism of *Molothrus ater* and the number of eggs in Icteridae species. Correlation between the parasitism of *M. ater* and the number of eggs per Icteridae species. On the left, white represents non-parasitized species, and black represents parasitized species. On the right, gray represents species that we don’t find reliable data, blue represents species that lay up to two eggs, green represents species that lay three or four eggs, black represents species that lay more than four eggs. Authors’ production.

**Fig 5.**
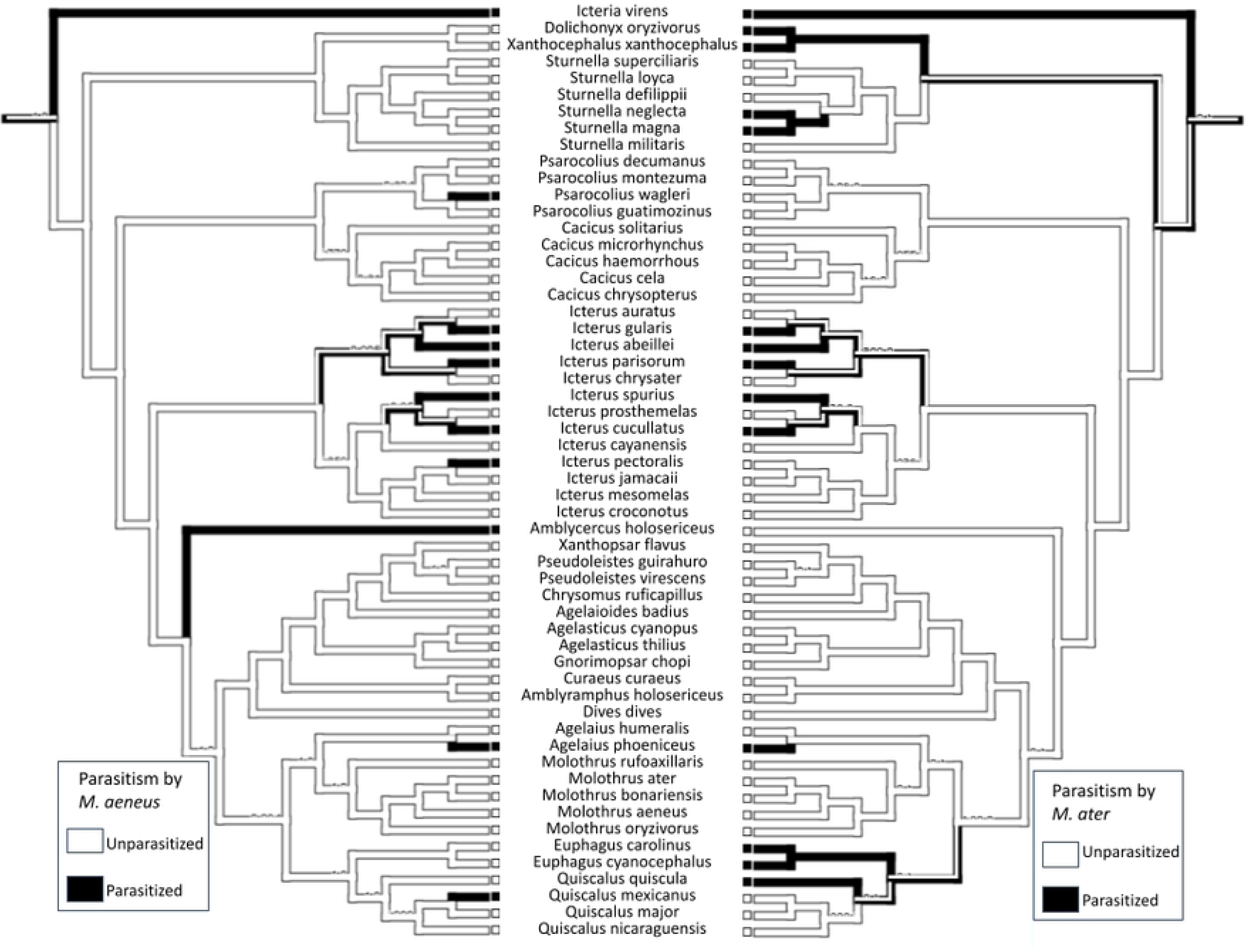
**Correlation between the parasitism of *Molothrus aeneus* and *Molothrus ater***. Correlation between the phylogenies of *M. aeneus* (left) and *M. ater* (right) in relation to the species they parasitize. Value 0, in white, represents non-parasitized species. Value 1, in black, represents parasitized species. Authors’ production.

#### Phylogenetic test for Bioregion

From a bioregional perspective, which reflects the real condition of the parasite-host interaction based on their co-occurrence, as species that do not co-occur cannot be subject to brood parasitism, we tested the correlations between each parasite and the bioregions in which they were present. Among them, only 3 out of 48 correlations were significant, all of them in bioregion 9 (Supporting information: Table 3), in relation to the total parasitism by *M. aeneus*, at points 3 (p = 0.0078), 4 (p = 0.03125), and 5 (p = 0.0156).

### Number of species parasitized by *Molothrus* spp. over evolution

Among the five species of *Molothrus*, we found a progression in the number of parasitized Icteridae species throughout the clade. *M. rufoaxillaris* parasitizes five species, *M. oryzivorus* parasitizes six, *M. aeneus* parasitizes eleven, *M. ater* parasitizes fourteen, and *M. bonariensis* parasitizes twenty-two species (Fig 6).

**Fig 6.**
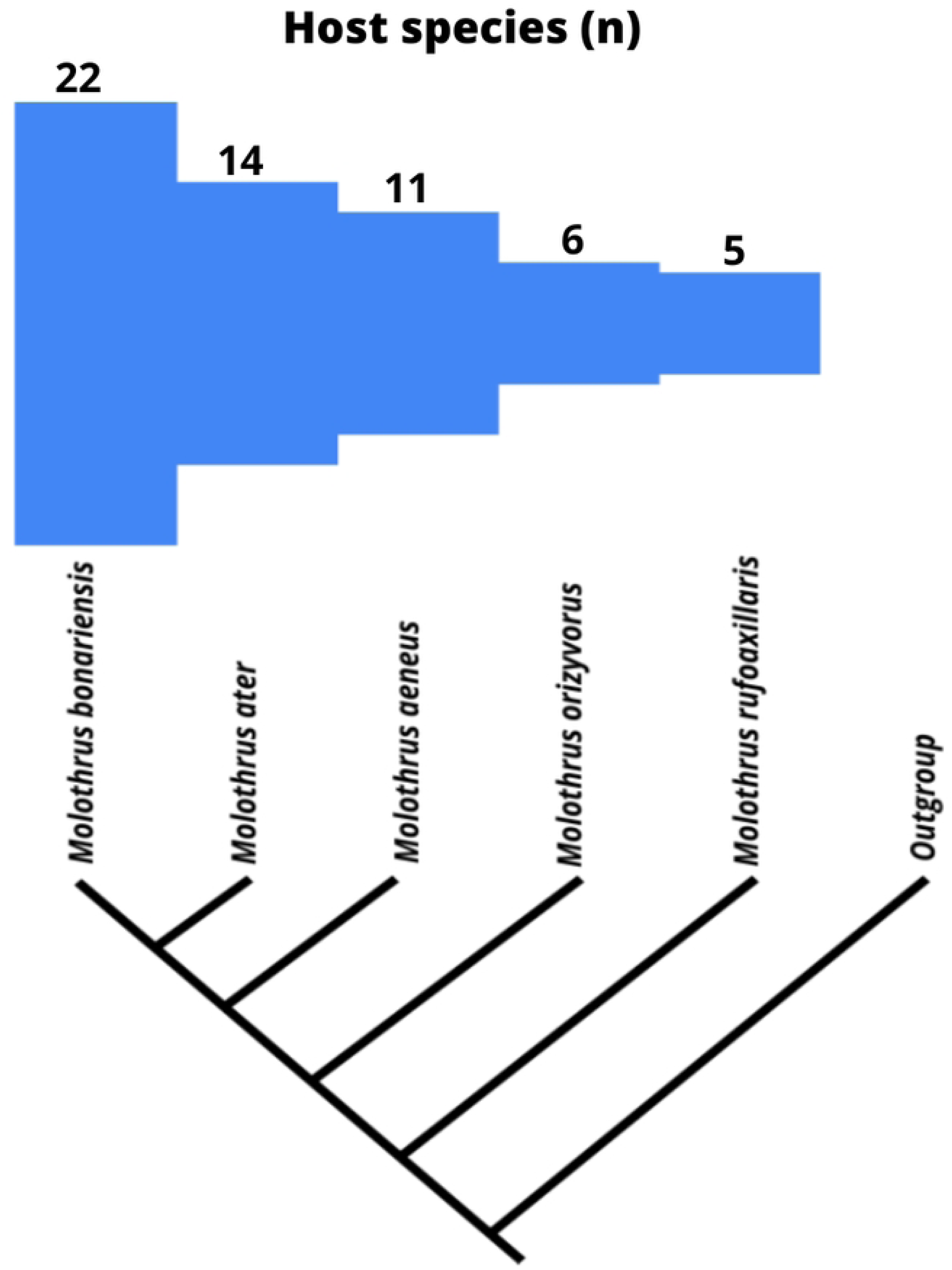
Cladogram showing the number of hosts of *Molothrus* spp. Cladogram depicting the phylogeny of *Molothrus* spp. and the number of icterid species they parasitize. Produced by the author’s based on LANYON (1992).

## Discussion

In our first hypothesis regarding the correlation between the different nest formats, diet, parental care and number of eggs of the hosts in relation to the brood parasites, we observed interesting results for nest types, as there was a noticeable preference of *M. oryzivorus* for nests with closed corridors. Regarding diet, no specialized selection by the parasites was identified, which refutes the relevance of this characteristic state in the parasite’s choice. In the case of the number of eggs, only one species, *M. ater*, showed a marginal result. There was also no proximal data regarding parental care.

Returning to the nest types, the choice of the host may not be solely determined by its nest type, as other species within the genera *Psarocolius* and *Cacicus* have closed nests with corridors but are not parasitized. It is plausible to consider that other previously studied characteristics are more fundamental factors in the structural condition of nests, with the prevention of nest parasitism being a secondary characteristic. This is similar to the case mentioned earlier regarding weaverbirds constructing closed and tubular nests to avoid predation and consequently, parasitism [2, 17].

It is correct to state that, according to the study conducted by Morelli, Benedetti, and Pape Møller (2020) on the diet of cuckoos, a dietary specialization was observed among nest-parasitic cuckoos compared to non-parasitic ones. However, in our study, there was no correlation found between diet and parasitism in Molothrus, indicating that this characteristic did not show an evolutionary trend in this case. A plausible explanation for this characteristic not interfering with *Molothrus* could be attributed to the diverse diet provided to bird nestlings, such as the gray-partridge (*Perdix perdix*), which receive invertebrate food from its parents in the first few weeks of life, while adults have a majoritarily plant-based diet [45–47].

Furthermore, the results did not allow us to make this statement because the factor “nest” may be associated with another characteristic, such as the number of eggs or parental care, which yielded negative results but were associated with the same species (*Psarocolius* and *Cacicus*). This could be a possible example of a “confounding variable,” as mentioned in the book by KREBS and DAVIES (1996), where a present characteristic is not necessarily linked to the study factor but to another selective pressure outside our focus. The same could apply to closed nests. Therefore, closed nests may prevent both parasitism and predation, as mentioned in the example by DAVIES, 2000, both being potential factors of evolutionary selection.

Our second hypothesis addressed an evolutionary trend in which increasingly closed nests throughout evolution would serve to prevent parasitism. When we analyzed the distribution of nest types on the cladogram (Fig 2), we observed that there was no linearity in the evolution of nest types. The open type (open) was the most basal, and the other types appeared independently throughout evolution. Thus, a selective pressure exerted by *Molothrus* did not occur, refuting this hypothesis. Nevertheless, we still see an indication that species constructing closed nests continue to be parasitized.

We should take into consideration studies on nest structure in passerine birds that show that nests of this order are open, but their common ancestor had closed nests [49–51]. This means that closed nests would not be a phylogenetic novelty but rather a feature that was lost and reappeared. Similar to the study by MEDINA et al. (2022), we believe that further research on nest types related to other factors still needs to be conducted for a better understanding of the evolutionary factors that select nest types.

In general, the genus *Molothrus* did not confirm the hypothetical preferences. This leads us to think that other factors that have been studied may have a greater influence. For example, there are studies that demonstrate factors such as morphological differences in eggs [9]; thicker eggshells in cases of parasites that puncture eggs [10]; size of the offspring in terms of feeding priority [53], and vocal mimicry by parasites imitating the begging calls of host offspring [1, 54].

Our third hypothesis concerns the increase in the number of species parasitized by *Molothrus* throughout its evolution. The results confirm the hypothesis by observing a quantitative progression in the number of parasitized species along the cladogram and indicate the success of the brood parasitic strategy over evolutionary time (Fig 6). This finding corroborates the phylogenetic proposal by LANYON (1992). However, it is important to identify some differences between this study and ours. In LANYON’s (1992) work, the available data about records of host species was probability smaller, since in our analysis, *M. rufoaxillaris* totaled five hosts in the Icteridae family, while LANYON (1992) mentioned only one. Another important point to highlight is that this increasing trend in the number of icterid hosts should be considered at each ancestral node of the cladogram in Figure 6. For evolutionary purposes, there is no difference between the 14 hosts of *M. ater* and the 22 hosts of *M. bonariensis*, as they are sister species with the same time of origin from the exclusive common ancestor. However, there has been an inversion of the species with the highest absolute number of hosts, not only in our analyses but also in the general spectrum [23].

We must emphasize that this strategy cannot be maximally efficient, as achieving complete parasitism of all nests would result in a lack of available hosts for the reproduction of the parasites. These results reveal a trend towards balanced evolution, without one species completely prevailing over the other, resembling an evolutionary arms race [5]. We can establish an analogy with the predator-prey relationship, where the total victory of predators would lead to the extinction of prey and, in turn, possibly the extinction of predators dependent on them. Similarly, if prey were always victorious in every encounter with predators, there would be no surviving predators as they would die of starvation. In the case of brood parasitism, it is likely that there is an alternation in the advantages obtained by hosts and brood parasites under different circumstances, resulting in different levels of success for each group.

An alternative testable hypothesis that could explain this possible equilibrium is that the number of parasites should be lower than that of hosts. However, parasites that exploit more host species might be more abundant and/or have larger geographic ranges compared to their congeners.

Finally, our last hypothesis addresses the correlation between *Molothrus* species sharing common hosts. In the phylogeny, we can observe a large number of Icteridae species parasitized by both *Molothrus ater* and *Molothrus aeneus* (Fig 4). In this case, the hypothesis was confirmed, and it is possible to consider it as an example of evolutionary convergence [34, 44], as they are not sister species, meaning they are not the closest relatives, but they exploit similar sets of hosts. Additionally, their geographic ranges have little overlap (IUCN). These narrow co-occurrence ranges provide an excellent opportunity to test whether they compete for host species. If we find a dispersed phylogenetic structure in sympatric Icteridae communities and clustered or null structure outside these sympatric zones, it may indicate competition for some resource in the areas of co-occurrence [34, 44].

It is worth emphasizing that our analyses have a taxonomic focus on a single family (Icteridae) within a hypothetical evolutionary scenario. If other species were added or removed from the regional pool, it would change the phylogeny and the regional pool itself, which could potentially alter the results and conclusions.

## Conclusion

It is possible to conclude that more open nests are indeed a more basal phylogenetic condition for the family. The presence of closed nest structures is not associated with a lower number of parasites, and further studies need to be conducted for a better understanding of this topic. The type of diet and parental care are not limiting factors for parasitism in a hypothetical scenario. The number of eggs shows a marginal correlation only for *Molothrus ater*, and the strategy of brood parasitism has persisted throughout evolution, seemingly increasing the number of exploited hosts. Additionally, conducting new studies on how each characteristic evolved based on all bird species would be interesting, as we were unable to compare our phylogenetic data with other similar contextual studies. Furthermore, testing these hypotheses in other families parasitized by *Molothrus* spp. can both corroborate and bring different perspectives to the results presented here.

## Acknowledgments

We would like to express our gratitude to the Federal University of Alfenas, particularly the Institute of Natural Sciences, for providing the opportunity to conduct our research. We would also like to thank Lucas Silva Azeredo for providing us with the initial idea and material for this work.

